# Reprogramming brain immunosurveillance with engineered cytokines

**DOI:** 10.1101/2022.06.21.497082

**Authors:** Anthony Tabet, Yash Agarwal, Jordan Stinson, Caroline Apra, Veronica Will, Marie Manthey, Noor Momin, Allison Sheen, Mitchell Murdock, Luciano Santollani, Li-Huei Tsai, Isaac Chiu, Sean Lawler, Darrell J. Irvine, K. Dane Wittrup, Polina Anikeeva

**Affiliations:** Department of Chemical Engineering, Massachusetts Institute of Technology, Cambridge, MA; McGovern Institute for Brain Research, Massachusetts Institute of Technology, Cambridge, MA; Koch Institute for Integrative Cancer Research, Massachusetts Institute of Technology, Cambridge, MA; Research Laboratory of Electronics, Massachusetts Institute of Technology, Cambridge MA; Department of Biological Engineering, Massachusetts Institute of Technology, Cambridge, MA; Sorbonne Universite, Paris, France; Department of Brain & Cognitive Sciences, Massachusetts Institute of Technology, Cambridge, MA; Picower Institute for Learning & Memory, Massachusetts Institute of Technology, Cambridge, MA; Broad Institute of MIT & Harvard, Cambridge, MA; Department of Immunology, Harvard Medical School, Boston, MA; Dept of Pathology & Laboratory Medicine, Legorreta Cancer Center, Brown University, Providence, RI; Department of Materials Science & Engineering, Massachusetts Institute of Technology, Cambridge MA; Ragon Institute of Massachusetts General Hospital, Massachusetts Institute of Technology and Harvard University, Cambridge, MA; Consortium for HIV/AIDS Vaccine Development, The Scripps Research Institute, La Jolla, CA; Howard Hughes Medical Institute, Chevy Chase, MD

## Abstract

Immune surveillance of the brain is regulated by resident non-neuronal cells and the blood-brain barrier.^1^ Dys-regulation of immunosurveillance is a hallmark feature of several diseases^2–5^ including brain tumors^6^ that interact with and rely heavily on immune cells,^7^ suggesting that disrupting the neuroimmunology of tumors could slow their progression. Yet few tools are available to control brain immunology *in vivo* with local precision, and fewer yet are used for therapeutic intervention. ^2^ Here, we propose engineered cytokines as a neuroimmune-modulation platform. We demonstrate that the residence time of cytokines in the brain can be tuned by binding them to the extracellular matrix or synthetic scaffolds. We then show that the aluminum hydroxide adjuvant (alum) is retained in the brain >2 weeks. Tethering of inflammatory cytokines such as interleukins (IL) 2 and 12 to alum yields extended neuroinflammation and brain immunosurveillance after intracranial administration, while avoiding systemic toxicity. In mouse models of both immunologically hot and cold brain tumors, the intracranial deposition of alum-tethered cytokines causes significant delay in tumor progression. RNA profiling reveals that engineered cytokines engage both innate and adaptive immunity in the brain. These findings suggest that engineered cytokines can reprogram brain immunosurveillance, informing the development of future therapies for neuroimmune diseases.

---

Classical notions of brain immune privilege and isolation are being challenged as increasing evidence reveals mechanisms of brain immunosurveillance by peripheral immune cells and antigen drainage through meningeal lymphatics. ^2,3,8,9^ Recent studies demonstrated that enhancing lym-phatic drainage through lymphangiogenic factor VEGF-C can improve monoclonal antibody therapy targeting amyloid beta in Alzheimer’s disease ^5^ or immune checkpoint blockade for brain cancer immunotherapy.^6^ Immunotherapy against established brain tumors could also be achieved with patientspecific chimeric antigen receptor (CAR) T cells,^10^ and in a recent clinical trial, CAR T cell immunotherapy was shown to improve clinical outcomes for children diagnosed with lethal pediatric gliomas.^11^ These elegant experiments highlight the utility of immune modulation and immunotherapy for treating central nervous system (CNS) diseases, and raise the question of whether other immuno-modulatory technologies such as engineered proteins can be adopted for CNS immunotherapy.

Interleukins (IL) 2 and 12 are inflammatory cytokines that polarize immune cells to mount a potent anti-cancer response.^12–15^ It was recently shown that intratumoral IL-12 can enhance CAR T cell immunotherapy in the brain, ^16^ and that IL-12 synergizes with checkpoint blockade.^17^ Despite their therapeutic promise, clinical translation of inflammatory cytokines is impeded by their systemic toxicity stemming from protein activity away from the tumor.^18–21^ In rodent models of peripheral cancers, cytokines fused to substrates with high residence times in tumors have been shown to trigger immune infiltration and activation while eliminating toxicity associated with systemic administration. ^20,22,23^ However, in contrast to the periphery, the ability of engineered cytokines to be retained in the brain and reprogram neuroinflammation is unknown given the putative immune-privileged status of this organ. Here, we test the hypothesis that extending the residence time of cytokines in the brain can similarly enhance immune rejection of established brain tumors and demonstrate anchored cytokines as a promising platform for neuroimmune engineering.

Anchoring of cytokines in peripheral tumors has been previously achieved by fusing them to proteins that bind to collagen in the extracellular matrix (ECM) or to synthetic scaffolds. To explore the utility of these two strategies in brain tissue, we chose the collagen-binding protein lumican as a means to anchor cytokines to the brain ECM, while aluminum hydroxide (alum) particles were chosen as a synthetic scaffold for binding cytokines fused to a phosphorylated alum-binding peptide (ABP, (Fig. 1a). Both of these anchoring strategies have been previously shown effective in eliciting immune activation in flank tumors.^20,22^ To assess the cytokine retention in the brain in the two anchoring scenarios, we first developed a simple finite element model of mass transport in the brain interstitium (Fig. 1b, Supplementary Information). Our model adopts features from recent brain transport simulations ^24^ into a proton mass transport model we recently reported.^25^ To capture the consequences of molecular weight on protein permeability, we rescaled a permeability model recently used to study protein transport in and out of flank tumors^23,26–28^

**Figure 1.**
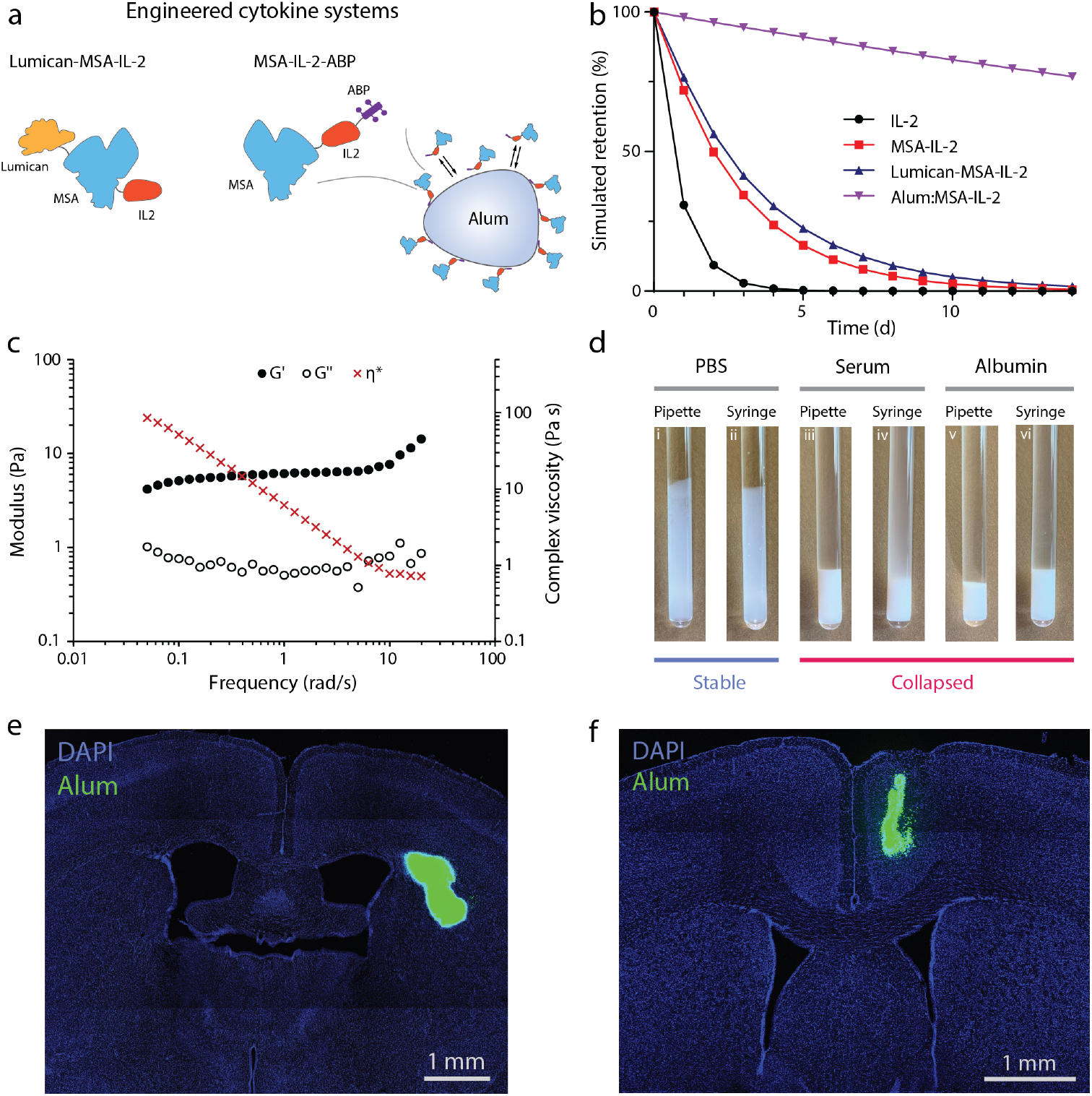
Alum is a weak particle suspension that collapses into a depot in the presence of serum and is retained in the brain for weeks. (a) Graphical illustration of the two engineered cytokine systems used to tune brain immunosurveillance. (b) Retention of various protein systems evaluated using a diffusion-convection-reaction transport model. (c) Frequency sweep done with oscillatory rheology showing the dynamic modulus of alum is on the order of 1-10 Pa. (d) Neither shear/extensional forces via syringe (compared to gentle pipetting) nor temperature trigger the collapse of alum into an agglomerated depot. Serum, or albumin at serum concentrations, is sufficient to cause alum to become unstable and collapse. (e-f) Alum is retained in the brain for at least 2 wks whether injected in the striatum or in the pre-motor cortex (M2).

The model predicts that lumican, which binds to collagen IV, fused to a cytokine of interest (IL-2 chosen as an example) increases its residence time in the brain compared to IL-2 alone or mouse serum albumin (MSA) fused to IL-2 (MSA-IL-2) (Fig. 1b). Since the abundance of collagen relative to other biomacromolecules is lower in the brain compared to many tissues in the periphery, the retention may not be as prolonged compared to flank tumors. In contrast, our model predicts that alum-tethered proteins would remain in brain tissue for over two weeks (77% at 14 d, Fig. 1b), suggesting its applications in longitudinal studies *in vivo*. Consistent with our model, intracranial administration of lumican-MSA-IL-2 into the striatum (Fig. S1) of immune-competent C57BL/6J mice invoked a local influx of CD3+ cells at 1 week (wk) but not 2 wks following delivery (Fig. S2). For mice injected with alum-tethered cytokines, the presence of CD3+ cells was found even after 2 weeks following delivery (Fig. S3), demonstrating that the duration of neuroinflammation can be predicted and tuned by changing the residence time of cytokines.

Being a common vaccine adjuvant, alum has been safely used in the clinic for nearly a century. ^20^ However, the effects of intracranially administered alum remain poorly characterized. The prior work in peripheral tissues ^20,22^ that informed our finite element model and our experimental observations of inflammatory response in the brain suggest that this material can remain in biological environments for extended periods of time. Yet the material properties of alum underlying this tissue retention remain unclear.

While handling alum, we observed the material does not hold its shape. Dynamic oscillatory rheology measurements reveal alum to be a weak gel with a dynamic modulus 0.01 kPa at room temperature (Fig. 1c), which is orders of magnitude lower than the moduli of biocompatible hydrogels commonly used for drug delivery (0.1-100 kPa) as well as the modulus of the brain tissue itself (1 kPa) ^29,30^ Given that these rheological properties of alum do not immediately suggest that it forms a tightly crosslinked depot *in vivo*, we investigated whether a temperature rise to 37°C, exposure to shear/extensional forces through injection, or the addition of serum (Fig. 1d-e) could change its properties. Neither temperature nor shear/extensional forces triggered macroscopic changes in alum suspensions, whereas the addition of serum proteins induced rapid alum precipitation (Fig. 1d). We then examined the ability of alum to collapse into such a depot in the brain by tagging the alum with a fluorescently labeled phosphoserine peptide and imaging brain slices 2 wks after injection (Fig. 1e-f). Interestingly, alum persists in the brain even 2 wks after injection in both the striatum (Fig. 1e) and pre-motor cortex (M2) (Fig. 1f). The retention of alum in the brain supports its utility in facilitating protein retention and a durable immune response in the context of brain tumors.

To investigate the therapeutic potential of alum-bound engineered cytokines, we inoculated immunologically infiltrated ‘hot’ orthotopic glioblastoma (GL261) or immunologically suppressive ‘cold’ syngeneic melanoma (B16F10) in the striatum of immune-competent mice and treated intratumorally with two cytokines, MSA-IL-2 (MSA used to enhance expression) and IL-12. Both cytokines were tethered to alum particles (Fig. 2a) by introducing a C-terminal alum-binding phosphoserine moiety (ABP). These phosphoserine affinity tags provide stable binding of proteins to alum *via* ligand exchange reactions of the phosphate groups with surface aluminum hydroxide groups.^20^ MSA-IL-2-ABP and IL-12-ABP both bind to alum, and binding of one does not sterically exclude the other from adsorbing (Fig. 2b-c, S4). In established GL261 glioblastomas (GBs), treatment with 4 *μ*g of MSA-IL-2 and 2 *μ*g of IL-12 tethered to 10 *μ*g of alum resulted in a substantial survival benefit over control saline, alum, or unanchored MSA-IL-2 and IL-12 treatments (Fig. 2d-f). Unanchored cytokines evoked systemic toxicity that caused 4 of 8 mice to reach euthanasia criteria 3 days following treatment (Fig. 2e). To treat the more aggressive B16F10 tumors, we increased the dosage to 10 *μ*g of MSA-IL-2 and 5 *μ*g of IL-12 tethered to 12.5 *μ*g of alum. To more closely match previously established treatment paradigms for B16F10 peripheral melanomas, ^20,22,31^ the tumor-targeting antibody TA99 and immune checkpoint blockade (anti-PD-1 monoclonal antibodies; aPD1) were also deployed intraperitoneally (i.p.). At these doses, saline or alum alone had poor survival outcomes (Fig. 2g-i). This dose of unanchored cytokines resulted in potent systemic toxicity, and all mice in that treatment group reached the euthanasia criteria 3 days after treatment. By contrast, cytokines tethered to alum did not elicit such pronounced systemic toxicity and resulted in significant survival benefit over the control groups (Fig. 2i). This therapeutic response was not specific to the striatum. We observed that alum is retained in both deep and more superficial cortical brain regions (Fig. 1e-f). When we inoculated melanoma tumors in the pre-motor cortex and treated mice 4 days later, we also observed a survival benefit (Fig. S5) with significant differences in locomotor behavior between treated and untreated animals (Fig. S6). The ability of the alum depot to collapse near where it was injected enables spatial specificity of cytokine localization for brain cancer immunotherapy.

**Figure 2.**
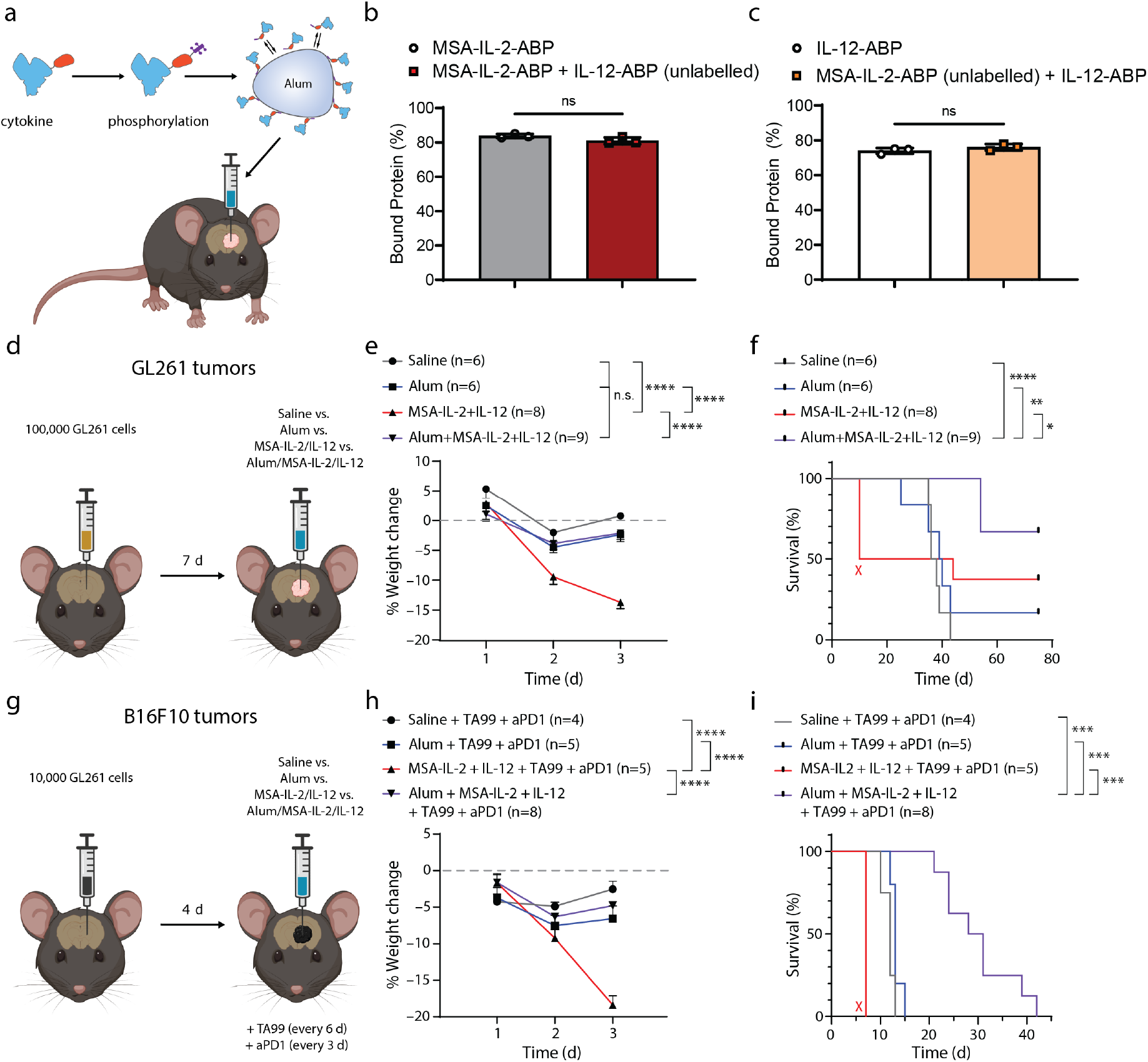
Alum-tethered cytokines can safely improve immunosurveillance against established brain tumors *in vivo*. (a) Graphic illustration demonstrating the strategy used to generate phosphorylated cytokines that adsorb to alum. (b) *In vitro* adsorption assay at 1 h demonstrating that MSA-IL-2 and IL-12 fused to the alum-binding phosphoserine moiety ABP can bind to alum at the same time without displacing the other. (c) Tumor inoculation and treatment strategy using orthotopic GL261 gliomas in the striatum of immune-competent C57BL/6J mice. (d) Weight loss data after treatment, demonstrating that intracranially injected MSA-IL-2 and IL-12 cause significant weight loss, whereas alum-tethered cytokines do not. An ordinary one-way ANOVA statistical analysis was conducted on day 3 weights. ****p<0.0001. (e) Alum-tethered cytokines impart a significant survival-benefit against orthotopic gliomas compared to all other groups. Statistical analysis was conducted with a Gehan-Breslow-Wilcoxon test. *p=0.0496. **p=0.0031. ****p<0.0001. (f) Tumor inoculation and treatment strategy using syngeneic B16F10 melanomas in the striatum of immune-competent C57BL/6J mice. (g) Weight loss data after treatment against B16F10 tumors, which included the tumor-targeting antibody TA99 and aPD1. Untethered cytokines in this paradigm cause significant weight loss; all mice reach euthanasia critera 3 d after treatment. An ordinary one-way ANOVA statistical analysis was conducted on day 3 weights. ****p<0.0001. (h) Alum-tethered cytokines impart a significant survival benefit compared to all other groups. Statistical analysis was conducted with a Gehan-Breslow-Wilcoxon test. Group 1 vs. Group 4 ***p=0.0005, Group 2 vs. Group 4 ***p=0.0004. Group 3 vs. Group 4 ***p=0.0005.

The ability of the alum depot to be retained near the injection site enables spatial resolution of cytokine localization for brain immunotherapy. Outside the CNS, IL-2 and IL-12 therapies rely on T cells.^20,22^ IL-2 and IL-12 promote T cell expansion and differentiation towards effector phenotypes, respectively.^32^ IL-12 also has the ability to directly engage with innate immune cells such as natural killer (NK) cells. As a vaccine adjuvant, alum triggers an inflammatory response^33^ that enhances trafficking to the depot, with subsequent antigen uptake and migration to the lymph node by infiltrating cells. Surprisingly, 2 wks after an alum-only injection in the brain, CD3+ cells could be found near the depot in both the striatum and the pre-motor cortex (Fig. 3a). This was surprising considering there were no tumors, no neo-antigens, nor any exogenous cytokines. Given this property, we next examined whether alum caused large changes to brain immunosurveillance everywhere, or if this effect was more spatially restricted, focal effect. We dissected out the dura 2 wks after injection and stained for CD3+ cells near the craniotomy or the sinus (Fig. 3b-c). We did not observe a very large population of T cells in the dura, suggesting again the effect on T cell trafficking was more focal.

**Figure 3.**
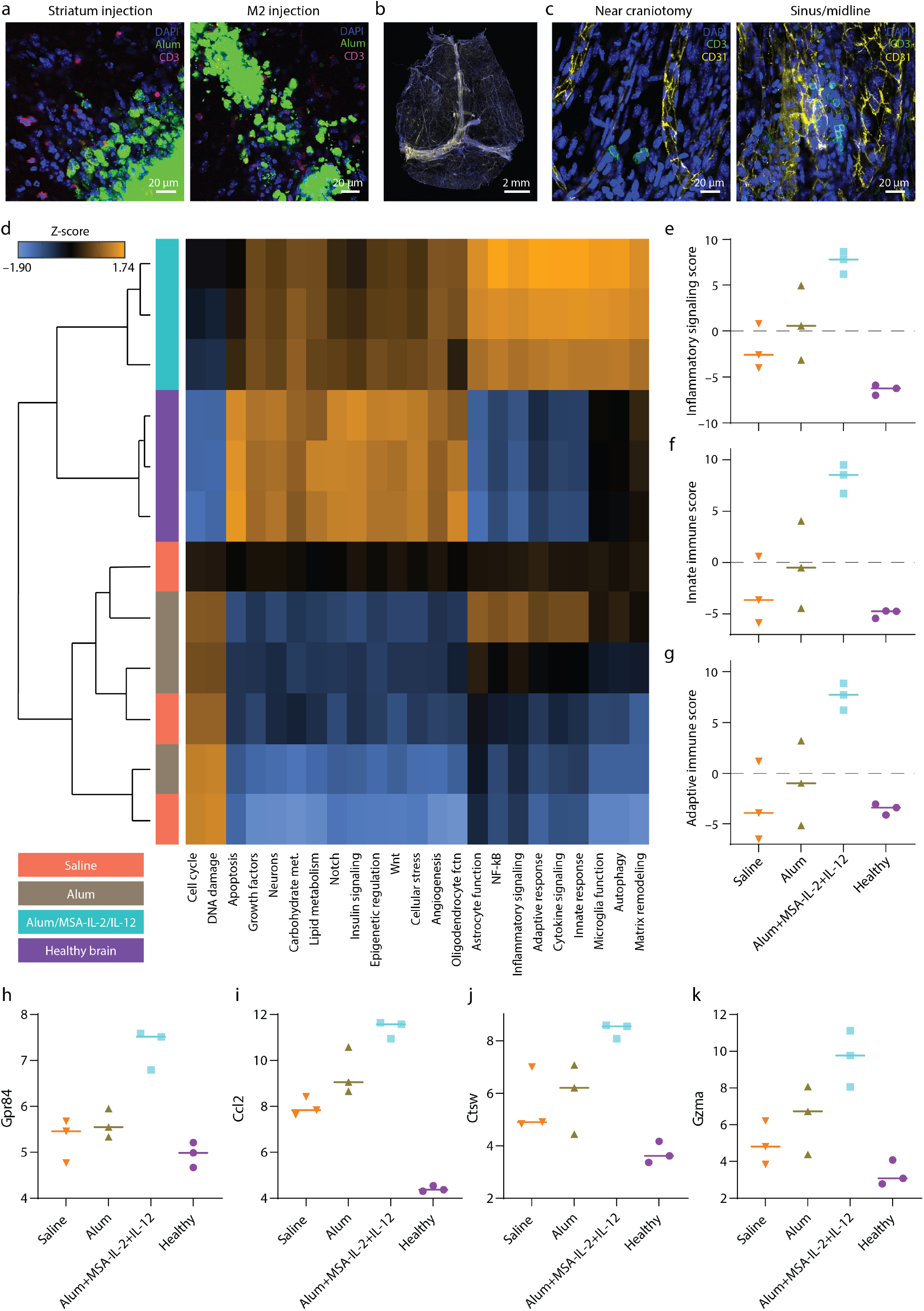
MSA-IL-2 and IL-12 tethered to alum engage innate and adaptive immune signaling cascades. (a) Confocal microscopy of striatum or pre-motor cortex (M2)-injected mice. CD3+ T cells traffic to the alum depot and persist 2 wks after injection. (b-c) We dissected out dura of these alum-injected mice and did not see a large influx of T cells near the craniotomy. (d-e) We performed RNA profiling using NanoString to obtain a more holistic immune profile of melanoma-bearing mice treated with saline, alum, or alum-tethered MSA-IL-2 and IL-12. Cytokine-treated mice exhibited a clear inflammatory profile. (f-g) alum/MSA-IL-2/IL-12 engage both innate and adaptive immune cascades, which are beneficial for anti-cancer immunity. (h-k) Alum/MSA-IL-2/IL-12 causes an increase of transcripts for (h) *Gp484*, associated with myeloid cells including microglia (i) *Ccl2*, a chemokine promoting migration of many cell types including DCs and T cells, (j) *Ctsw*, a marker of cytotoxic T and NK cells and (k) *Gzma*, a cytotoxic T and NK cell protease. Genes are plotted as log2 of normalized expression.

To further elucidate the neuroimmuno-modulatory effects of alum and alum-tethered cytokines, we quantified transcripts associated with cell proliferation, viability, and inflammation with RNA profiling. Mice were inoculated with B16F10 tumors in the striatum and treated with saline, alum only, or alum with MSA-IL-2 and IL-12 (all tumor-bearing mice also received TA99 and aPD1) 4 d later. RNA was extracted from brain tissue sections (Qiagen FFPE RNEasy) 10 d following tumor inoculation and analyzed through the nCounter(R) Mouse Neuroinflammation panel (Fig. 3, see Methods). In tumor-bearing mice injected with alum-tethered cytokines, a substantial increase in transcripts of genes associated with multiple inflammatory pathways was detected (Fig. 3d-e), suggesting that alum-tethered MSA-IL-2 and IL-12 engaged with innate and adaptive immune signaling cascades (Fig. 3f-g). This increase was not observed in healthy mice or tumor-bearing mice treated with saline or alum alone.

The tethered cytokines led to an increased count of *Gp484* transcripts, which are associated with myeloid cells such as microglia (Fig. 3h), as well as *Tnf* transcripts, which encode for the inflammatory cytokine produced by microglia, T cells, and NK cells (Fig. S7). We also observed upregulation of chemokine transcripts such as *Ccl2*, which promotes the infiltration of dendritic cells (DCs) and T cells (Fig. 3i). Furthermore, an upregulation of *Ctsw* (Fig. 3j) and *Gzma* (Fig. 3k), which are genes associated with cytotoxic T cell function, was observed in tethered cytokine-treated tumor-bearing mice. Overall, in subjects treated with alum-anchored cytokines, 342 genes are differentially expressed over saline-treated controls, compared to 3 genes being differentially expressed in subjects treated with alum alone (Fig. S8). Interestingly, the 3 differentially expressed genes were CCl2, Ccl7, and Tgm1, supporting the notion that alum itself exhibits inflammatory effects, though significantly lower than those associated with bound cytokines.

Interestingly, the brain tissue of tumor-bearing mice treated with alum-anchored cytokines was enriched for expression of genes commensurate with healthy brain function, including those involved in carbohydrate or lipid metabolism, which was more similar to brain tissue from naïve subject than the animals bearing tumors but treated with saline or alum alone (Fig. 3d). Similar trends were observed for genes marking neuron and oligodendrocyte viability (*Grm2, Myrf*, and *Tmem88b*, Fig. S9). Additionally, cells important to blood-brain barrier (BBB) maintenance and function were closer to a healthy brain in mice treated with anchored cytokines (Fig. S9). These findings consistent with observations of behavior (Fig. S6) suggest that alum-tethered cytokines preserve neuronal function and dynamics for greater periods of time at least in part by promoting immune surveillance of tumors.

In this work, we show that engineered cytokines can be used to reprogram the immune profile of the brain and trigger neuroinflammation for different periods of time (Fig. S10). We also show that alum-tethered cytokines impede the progression of established brain tumors. Investigating the material properties of alum revealed that alum forms a depot in the brain that persists for >2 wks. Intratumorally, intracranially injected alum-tethered MSA-IL-2 and IL-12 cause a potent anti-tumor immune response and engage both innate and adaptive immune mechanisms. Our findings indicate that engineered proteins can safely and locally alter the immune profile of the brain, empowering studies of neuroimmune interactions in health and disease.

## Acknowledgement

A.T., A.S., and L.S. thank the National Science Foundation Graduate Research Fellowship Program for funding support. A.T. thanks the Paul and Daisy Soros Fellowship for funding support. C.A. thanks the Ligue Contre le Cancer, the Fondation Bettencourt-Schueller, the Fondation Fyssen, the Philippe Foundation, the Institut Servier, the French Neurosurgical Society and the Assistance Publique-Hopitaux de Paris for funding support. P.A. thanks the support of the Dresselhaus Fund Award, the Department of Materials Science and Engineering Start-up fund, and the K. Lisa Yang and Hock E. Tan Center for Molecular Therapeutics. The authors acknowledge the Koch Institute Histology Core and the MIT BioMicro Center.

## References

(1) Sweeney, M. D.; Zhao, Z.; Montagne, A.; Nelson, A. R.; Zlokovic, B. V. Blood-Brain Barrier: From Physiology to Disease and Back. Physiological Reviews 2019, 99, 21–78, PMID: 30280653.

(2) Tabet, A.; Apra, C.; Stranahan, A. M.; Anikeeva, P. Changes in brain neuroimmunology following injury and disease. Frontiers in Integrative Neuroscience 2022,

(3) Louveau, A.; Smirnov, I.; Keyes, T. J.; Eccles, J. D.; Rouhani, S. J.; Peske, J. D.; Derecki, N. C.; Castle, D.; Mandell, J. W.; Lee, K. S.; Harris, T. H.; Kipnis, J. Structural and functional features of central nervous system lymphatic vessels. Nature 2015, 523.

(4) Da Mesquita, S. et al. Functional aspects of meningeal lymphatics in ageing and Alzheimer’s disease. Nature 2018, 560.

(5) Mesquita, S. D. et al. Meningeal lymphatics affect microglia responses and anti-A immunotherapy. Nature 2021, 593.

(6) Song, E.; Mao, T.; Dong, H.; Boisserand, L. S. B.; Antila, S.; Bosenberg, M.; Alitalo, K.; Thomas, J. L.; Iwasaki, A. VEGF-C-driven lymphatic drainage enables immunosurveillance of brain tumours. Nature 2020, 577.

(7) Cordell, E. C.; Alghamri, M. S.; Castro, M. G.; Gutmann, D. H. T lymphocytes as dynamic regulators of glioma pathobiology. Neuro-Oncology 2022, noac055.

(8) Louveau, A. et al. CNS lymphatic drainage and neuroinflammation are regulated by meningeal lymphatic vasculature. Nature Neuroscience 2018, 21.

(9) Louveau, A.; Harris, T. H.; Kipnis, J. Revisiting the Mechanisms of CNS Immune Privilege. Trends in Immunology 2015, 36.

(10) Mount, C. W.; Majzner, R. G.; Sundaresh, S.; Arnold, E. P.; Kadapakkam, M.; Haile, S.; Labanieh, L.; Hulleman, E.; Woo, P. J.; Rietberg, S. P.; Vogel, H.; Monje, M.; Mackall, C. L. Potent antitumor efficacy of anti-GD2 CAR T cells in H3-K27M+ diffuse midline gliomas letter. Nature Medicine 2018, 24, 572–579.

(11) Majzner, R. G. et al. GD2-CAR T cell therapy for H3K27M-mutated diffuse midline gliomas. Nature 2022,

(12) Briukhovetska, D.; Dörr, J.; Endres, S.; Libby, P.; Dinarello, C. A.; Kobold, S. Interleukins in cancer: from biology to therapy. Nature Reviews Cancer 2021, 21.

(13) Holder, P. G.; Lim, S. A.; Huang, C. S.; Sharma, P.; Dagdas, Y. S.; Bulutoglu, B.; Sockolosky, J. T. Engineering interferons and interleukins for cancer immunotherapy. Advanced Drug Delivery Reviews 2022, 182, 114112.

(14) Barrett, J. A.; Cai, H.; Miao, J.; Khare, P. D.; Gonzalez, P.; Dalsing-Hernandez, J.; Sharma, G.; Chan, T.; Cooper, L. J. N.; Lebel, F. Regulated intratumoral expression of IL-12 using a RheoSwitch Therapeutic System® (RTS®) gene switch as gene therapy for the treatment of glioma. Cancer Gene Therapy 2018, 25, 106–116.

(15) Parker, J. N.; Gillespie, G. Y.; Love, C. E.; Randall, S.; Whitley, R. J.; Markert, J. M. Engineered herpes simplex virus expressing IL-12 in the treatment of experimental murine brain tumors. Proceedings of the National Academy of Sciences 2000, 97, 2208–2213.

(16) Agliardi, G. et al. Intratumoral IL-12 delivery empowers CART cell immunotherapy in a pre-clinical model of glioblastoma. Nature Communications 2021, 12, 444.

(17) vom Berg, J.; Vrohlings, M.; Haller, S.; Haimovici, A.; Kulig, P.; Sledzinska, A.; Weller, M.; Becher, B. Intratumoral IL-12 combined with CTLA-4 blockade elicits T cell–mediated glioma rejection. Journal of Experimental Medicine 2013, 210, 2803–2811.

(18) Leonard, J. P.; Sherman, M. L.; Fisher, G. L.; Buchanan, L. J.; Larsen, G.; Atkins, M. B.; Sosman, J. A.; Dutcher, J. P.; Vogelzang, N. J.; Ryan, J. L. Effects of Single-Dose Interleukin-12 Exposure on Interleukin-12–Associated Toxicity and Interferon-Production. Blood 1997, 90, 2541–2548.

(19) Pfreundschuh, M. G.; Tilman Steinmetz, H.; Tüschen, R.; Schenk, V.; Diehl, V.; Schaadt, M. Phase I study of intratumoral application of recombinant human tumor necrosis factor. European Journal of Cancer and Clinical Oncology 1989, 25, 379–388.

(20) Agarwal, Y.; Milling, L. E.; Chang, J. Y. H.; Santollani, L.; Sheen, A.; Lutz, E. A.; Tabet, A.; Stinson, J.; Ni, K.; Rodrigues, K. A.; Moyer, T. J.; Melo, M. B.; Irvine, D. J.; Wittrup, K. D. Intratumourally injected alum-tethered cytokines elicit potent and safer local and systemic anticancer immunity. Nature Biomedical Engineering 2022, 6, 129–143.

(21) Merchant, R. E.; McVicar, D. W.; Merchant, L. H.; Young, H. F. Treatment of recurrent malignant glioma by repeated intracerebral injections of human recombinant interleukin-2 alone or in combination with systemic interferon-α. Results of a phase I clinical trial. Journal of Neuro-Oncology 1992, 12, 75–83.

(22) Momin, N.; Mehta, N. K.; Bennett, N. R.; Ma, L.; Palmeri, J. R.; Chinn, M. M.; Lutz, E. A.; Kang, B.; Irvine, D. J.; Spranger, S.; Wittrup, K. D. Anchoring of intratumorally administered cytokines to collagen safely potentiates systemic cancer immunotherapy. Science Translational Medicine 2019, 11.

(23) Momin, N.; Palmeri, J. R.; Lutz, E. A.; Jailkhani, N.; Mak, H.; Tabet, A.; Chinn, M. M.; Kang, B. H.; Spanoudaki, V.; Hynes, R. O.; Wittrup, K. D. Maximizing response to intratumoral immunotherapy in mice by tuning local retention. Nature Communications 2022, 13, 109.

(24) Ray, L.; Iliff, J. J.; Heys, J. J. Analysis of convective and diffusive transport in the brain interstitium. Fluids and Barriers of the CNS 2019, 16.

(25) Park, J.; Tabet, A.; Moon, J.; Chiang, P.-H.; Koehler, F.; Sahasrabudhe, A.; Anikeeva, P. Remotely Controlled Proton Generation for Neuromodulation. Nano Letters 2020, 20, 6535–6541.

(26) Schmidt, M. M.; Wittrup, K. D. A modeling analysis of the effects of molecular size and binding affinity on tumor targeting. Molecular Cancer Therapeutics 2009, 8.

(27) Thurber, G. M.; Wittrup, K. D. A mechanistic compartmental model for total antibody uptake in tumors. Journal of Theoretical Biology 2012, 314.

(28) Wittrup, K. D.; Thurber, G. M.; Schmidt, M. M.; Rhoden, J. J. Practical theoretic guidance for the design of tumor-targeting agents. Methods in Enzymology 2012, 503.

(29) Li, J.; Mooney, D. J. Designing hydrogels for controlled drug delivery. 2016, 1.

(30) Tabet, A.; Mommer, S.; Vigil, J. A.; Hallou, C.; Bulstrode, H.; Scherman, O. A.. Adv. Healthcare Mater. 2019, 7, 1827.

(31) Moynihan, K. D. et al. Eradication of large established tumors in mice by combination immunotherapy that engages innate and adaptive immune responses. Nature Medicine 2016, 22.

(32) Kieper, W. C.; Prlic, M.; Schmidt, C. S.; Mescher, M. F.; Jameson, S. C. IL-12 Enhances CD8 T Cell Homeostatic Expansion. The Journal of Immunology 2001, 166.

(33) Moyer, T. J. et al. Engineered immunogen binding to alum adjuvant enhances humoral immunity. Nature Medicine 2020, 26.

