## Supplementary Information for "Reprogramming brain immunosurveillance with engineered cytokines"

### S.1 Materials and Methods

#### Protein production

We previously published an extensive protocol on protein production and purification for the cytokines reported here[1]. Proteins used in this work were produced and purified identically as previously reported. Briefly, genes for ABP-containing MSA–IL2 and IL12, Fam20C (Horizon), and lumican-MSA-IL2 were transformed into Stellar Competent Cells (Takara) and purified using the NucleoBond Xtra Maxi EF endotoxin-free maxi-prep kit (Takara).

Plasmids were transfected (transiently) into HEK293F cells for protein production (1 mg total DNA L−1 cell culture) with polyethylenimine (2 mg L−1 cell culture) using Freestyle 293 Expression system (Gibco). Protein plasmid:Fam20C plasmid transfection mass ratio was restricted to 9:1. TA99 was purified using rProtein A Sepharose Fast Flow resin (Cytiva Life Sciences), as previously reported[1]. These his-tagged proteins from cell culture supernatants were purified using HisPur Ni-NTA metal affinity resin (Thermo Fisher Scientific). Phosphorylated proteins were further purified using anion exchange chromatography (see Agarwal et al[1] for an extended discussion on phosphorylated protein purification).

Before *in vivo* administration, proteins were buffer exchanged into tris-buffered saline (Sigma-Aldrich) using Amicon spin columns. Purified proteins were confirmed to have low endotoxin levels (<0.1 EU per dose) with Charles River Endosafe Nexgen-PTS.

**In vitro binding assay**

In vitro binding was determined as previously report [1]. Briefly, proteins were conjugated to AF647 with NHS chemistry (Invitrogen), mixed with alum, and rotated at room temperature for 20 min. Samples were then centrifuged at 10,000g for 10 min to form a pellet, and the supernatant was collected and replaced with 10% mouse serum-containing PBS. The tubes were then rotated at 37C. At 1 h, 1 d, and 4 d, samples were again centrifuged, and the supernatant was again replaced with 10% mouse serum. The fluorescence of the supernatants were analyzed using a Tecan Infinite M200 Pro plate reader. Results were normalized to no-alum samples.

**Animals**

In this study, 6 to 10-week-old female C57BL/6 (Jackson Laboratory) were used. Mice were housed at the MIT animal facility, and regular housing conditions (12 h L/D cycle, 22 °C, food and water *ad libitum*) were used. Animals were monitored daily for 4 days after surgeries and 2-3 times per week thereafter. The euthanasia criteria were 15% weight loss compared to day 0 weight, extensive loss of weight (20%) since last monitoring event, BCS index < 2, comatose or moribund state. All experiments were approved by the MIT Committee on Animal Care.

**Cells**

In this study, GL261 and B16F10 murine cells were used. Trypsin-EDTA (0.25%), and fetal bovine serum were purchased from Invitrogen. Dulbecco's Modified Eagle's Medium (DMEM, high glucose) was purchased from ATCC. Cells were passaged in DMEM with 10% FBS. For tumor inoculations, cells were washed with PBS, trypsinized, washed with DMEM and then with PBS, before being resuspended in either 100,000 cells / 3 $\mu$L for GL261 or 10,000 cells / 1 $\mu$L for B16F10. Cells were injected at 1 $\mu$L/min *in vivo*.

**Craniotomies**

All animal experiments were approved by the MIT Committee on Animal Care. Surgeries were performed on deeply anesthetized mice (isoflurane) and positioned in a stereotactic frame (David Kopf Instruments). 28 gauge Hamilton needles were used. Striatum injections occurred with the following coordinates: -0.5 mm anterior-posterior, 2.0 mm mediolateral, -2.7 mm deep to cranial surface. Pre-motor cortex injections occurred with the following coordinates: 1.0 mm anterior-posterior, 0.5 mm mediolateral, -1.25 mm deep to cranial surface. For all injections, the infusion was started at 0.2 mm deeper than the coordinate and the needle was slowly raised to the final position.

**Treatments**

Mice bearing GL261 tumors were treated on day 7 with 10 ug of alum, 4 ug of MSA-IL2, and 2 ug of IL12. Mice bearing B16F10 tumors were treated on day 4 with 12.5 ug of alum, 10 ug of MSA-IL2, and 5 ug of IL12, with the exception of mice with M2 B16F10 tumors, which were treated with the lower dose. Mice bearing B16F10 tumors were also treated with 100 ug of TA99 every 6 days for the duration of the study and 200 ug of aPD1 (clone 29F.1A12, Bio X Cell) every 3 days for the duration of the study. TA99 and aPD1 were administered with intraperitoneal injections.

### Behavioral experiments

In the open field behavioral assay, adult female mice (C57Bl/6, 8-10 weeks old at start of the experimental procedure) where used during the light phase of the light/dark circle, and were acclimated 1-2 hours to the behavior room as well as to the handling procedure for 3 consecutive days before the first experimental day.

On the days of testing, the mice were brought into the set up room 1-2 hours before testing. The behavior room was set up with consistent white light and minimal outside noise. The mice were placed into the center of a home-built box (30cm x 30cm) out of polyethylene sheets (McMaster) with a thin layer of bedding for 15 min of total recording time. Video recordings were done with a webcam (Logitech) and its respective software, and positioned above the center of the open field chamber. After recording, the mice were placed back into their homecage and the open field chamber was cleaned and laid out with fresh bedding.

The recorded video was imported into Ethovision IX software and the recorded time from minute 6 to minute 15 was analyzed for average distance traveled and average velocity. Each mouse from each group was tested on days 7, 10, 14, 18 relative to tumor inoculation in the open field.

### Finite element modeling

Mass transport was modeled with COMSOL Multiphysics finite element package (5.4). We built a semi-quantitative convection-diffusion-reaction transport model in water to compare the effects of molecular weight or substrate binding had on relative retention of IL2. The built-in properties of water were used. The initial condition protein concentration was placed on a 1.5 mm diameter circle at the center of a larger 10.0 mm circle. Cytokine initial condition was set to 0 outside the inner circle. We set an instantaneous consumption as the outermost boundary condition ($k_{surf}$ = $10^{13}$ m/s. In the collagen-binding models, a concentration of 112 nM was set as the brain lumican binding site initial condition everywhere in the simulation. This value was obtained by taking a hydroxyproline (HP)/binding site ratio[2] and rescaling to account for the HP content in collagen IV and the void fraction. To capture convective mass transport, we placed 16 arteries or venules within the initial protein bolus (Fig. S11), which another report recently used to model convection and diffusion in brain interstitium[3]. The arteries and venules diffusivity scaling was also used. The outer diameter of each was 0.09 mm, and the inner diameter was 0.03 mm. Using Darcy's Law, we set the boundary condition pressure as 0 on the first and third columns, and 4 mmHg on the second and fourth columns. Protein retention was calculated within the square around the 16 arterioles or venules (Fig. S11).

To capture the effects of molecular weight on protein permeability, we rescaled a permeability model recently used to study protein transport in and out of flank tumors[2, 4–6] by a factor of $71.8^{-1}$. This rescaled value was obtained by matching this permeability model to published brain permeability values of macromolecules[7] at various molecular weights (4, 20, 70 kDa). This permeability (P) was used as a sink in the 16 arterioles or venules (Fig. S11) for free luminca-MSA-IL2, MSA-IL2, or IL2 where the flux was equal to the permeability multiplied by the concentration. Proteins that were bound to collagen or alum were given a diffusivity and permeability of 0 as they were immobilized.

The sinks that changed protein concentration within the square bound (Fig. S11) were (1) permeability of protein across vessels and (2) convective/diffusive transport of protein away from the initial bolus. Lumican-MSA-IL2 could bind to a collagen binding site ($K_D$=100 nM)[8]. While not captured in this model, an advantage of collagen-binding proteins is that when they leave the tumor, they do not traffic to the blood in the way unbound MSA-IL2 does, but instead they are entrapped in other collagen-rich structures, such as lymph nodes[2]. Protein desorption from alum was set as irreversible, with $k_{off}$ set as $2.17*10^{-7}$ $s^{-1}$. This value was obtained by assuming a reaction:

$$P_{bound} \rightarrow P_{free}$$

$$\frac{dP_{bound}}{dt} = -k_{off}P_{bound}$$

We previously measured approximately 30% *in vivo* signal loss over 19 days (starting on day 5)[1], and this data was used to obtain $k_{off}$. The remaining parameters used in this model are as follows:

| Parameter | Value |
|---|---|
| Hydraulic conductivity K | 1.5001E-12 $m^3 \cdot s/kg$ |
| Tortuosity | 1.6 |
| Diffusivity of MSA-containing proteins | 6E-11 $m^2/s$ |
| Diffusivity of IL2 | 1E-10 $m^2/s$ |
| lumican $k_{on}$ | 10 $m^3/(s \cdot mol)$ |
| lumican $k_{off}$ | 0.001 1/s |
| Diffusivity of MSA-containing proteins | 6E-11 $m^2/s$ |
| Diffusivity of IL2 | 1E-10 $m^2/s$ |
| Permeability of IL2 | 2.13E-10 m/s |
| Permeability of MSA-IL2 | 5.35E-11 m/s |
| Permeability of lumican-MSA-IL2 | 4.48E-11 m/s |

**Fixation and brain confocal microscopy**

Animals were anesthetized with isoflurane, injected with fatal plus (100 mg/kg IP), and transcardially perfused with 50 mL of ice cold phosphate buffered saline (PBS) followed by 50 mL of ice cold 4% paraformaldehyde (PFA) in PBS. The brains were removed and post-fixed in 4% PFA at 4 °C. Confocal microscopy images were obtained using a laser scanning confocal microscope (FluoView FV1000, Olympus). Brain slices (50 microns) were dried and mounted using Fluoromount. 4x, 10x, 20x, and 60x objectives were used for imaging. Slices were blocked in PBSTA (0.3% triton-X, 3% bovine serum albumin) for 1 h, antibody stained overnight at room temperature and 100 RPM in PBSTA, washed with PBS, DAPI stained for 30 min, washed again, and mounted. We used a AF594-conjugated CD3 antibody from Biolegend (100240). To avoid wound-healing artifacts in the CD3 quantification experiments, CD3+ cells were quantified at bregma above and below the ventricle in the hemisphere where the injection took place at 1 or 2 weeks after injection of saline or inflammatory cytokines.

**Dura dissections**

Whole mounts of the dural meninges were prepared as follows. Following perfusion, skull caps were removed, then placed in 4% paraformaldeyhyde at 4ºC for 12 hours. The dural meninges (dura mater and arachnoid) were peeled from the skull cap under a dissecting microscope using Dumont forceps (Fine Science Tools) then placed in a 24-well plate (VWR 10861-558) with phosphate buffered saline (PBS) for immunohistochemistry. First, meninges were washed with PBS for 10 minutes, permeabilized with 0.3% Triton X-100 in PBS for 10 minutes, underwent blocking (5% Normal Donkey Serum and 0.3% Triton X-100 in PBS) for 1 hour at room temperature, and immunostained with the primary antibodies in blocking solution overnight at room temperature. For primary antibodies, we used Armenian Hamster-CD31 (Sigma-Aldrich MAB1398Z) and CD3 Rat anti-Mouse Clone 17A2 (eBioscience 14-0032-85) (dilution of 1:500 for each antibody). Following three five-minute washes with blocking buffer, we added appropriate secondary antibodies for 2 hours at room temperature, then washed with PBS five times for five minutes each. On the penultimate wash we used 1:1000 Hoechst (Thermo Fisher Scientific, H3570). Tissue was mounted on SuperFrost slides and sealed with Prolong Gold mounting medium (Thermo Fisher Scientific, P36930).

**RNA Profiling**

Animals were anesthetized with isoflurane, injected with fatal plus (100 mg/kg IP), and transcardially perfused with 50 mL of ice cold phosphate buffered saline (PBS) followed by 50 mL of ice cold 4% paraformaldehyde (PFA) in PBS. The brains were removed and post-fixed in 4% PFA at 4 °C. In line with Nanostring recommendations for sample preparation, brains were fixed for 18 hours in 4% PFA prior to tissue processing and paraffin-embedding. For each sample, embedded tissue blocks were faced off prior to collecting three 10um scrolls per sample. Excess paraffin was trimmed prior to RNA extraction using the Qiagen RNEasy FFPE kit. RNA quantity and quality were assessed using Fragment Analyzer (Agilent). 100ng of sample RNA >200nt was hybridized for 20 hours to the Nanostring nCounter Mouse Neuroinflammation CodeSet and then loaded onto the sample cartridge using Nanostring Max/Flex system. The cartridge was scanned at the highest quality and data analysis and normalization was performed in nSolver v4.0. Assessment of differentially expressed genes, pathway enrichment, and cell type profiling was performed in Nanostring nSolver Advanced Analysis v2.0.

**Rheology**

Rheological sweeps were performed on an AR-G2 Rheometer (TA Instruments, New Castle, DE, USA) with a 20 mm parallel plate geometry at 20°C. Hydrogel samples were loaded onto the rheometer with a 1000 µm loading gap.

**Statistics**

GraphPad Prism (9.2.0) was used for all statistical analyses. The specific statistical tests used for each data are written in the figure captions where the data appears.

### Safety

No unexpected or unusually high safety hazards were encountered in the experiments reported here.

### S.2 Supporting Figures

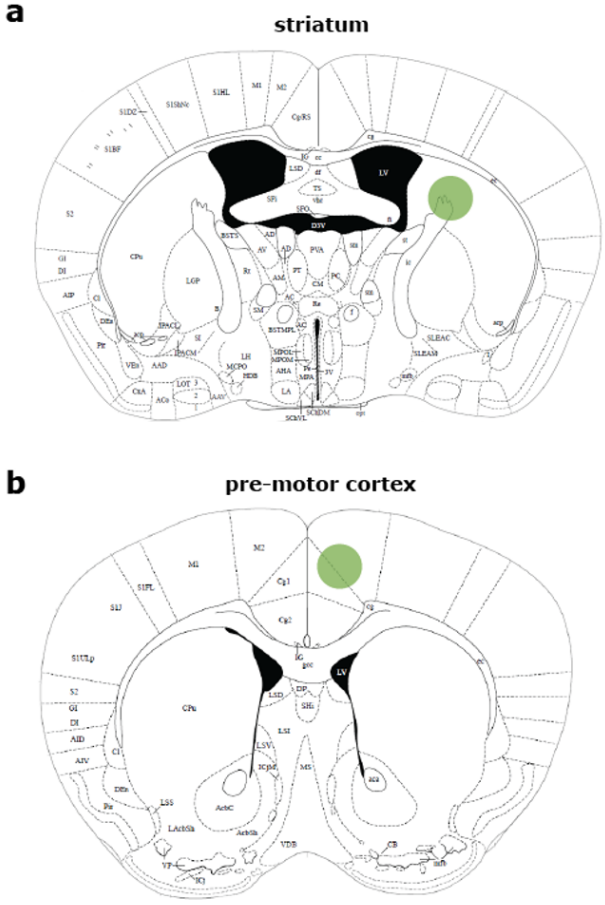


**Figure S1**: (a) Striatum and (b) pre-motor cortex are the brain regions targeted in this work. Images were adopted from the Mouse Brain Atlas[9].

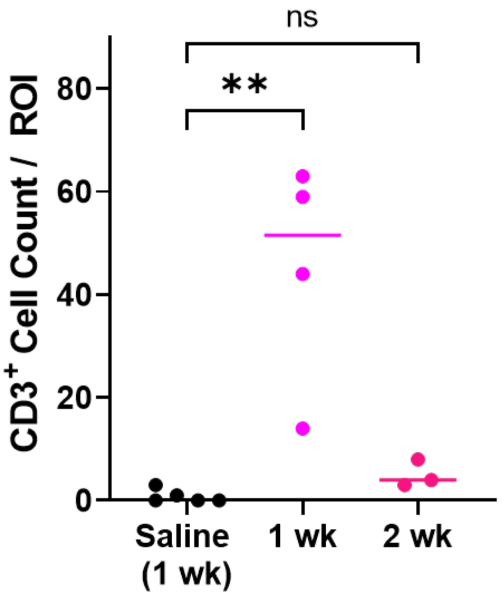


**Figure S2**: Quantification of CD3+ cells after lumican-MSA-IL2 administration into the striatum. A one-way ANOVA was used for statistical analysis. **p=0.0013.

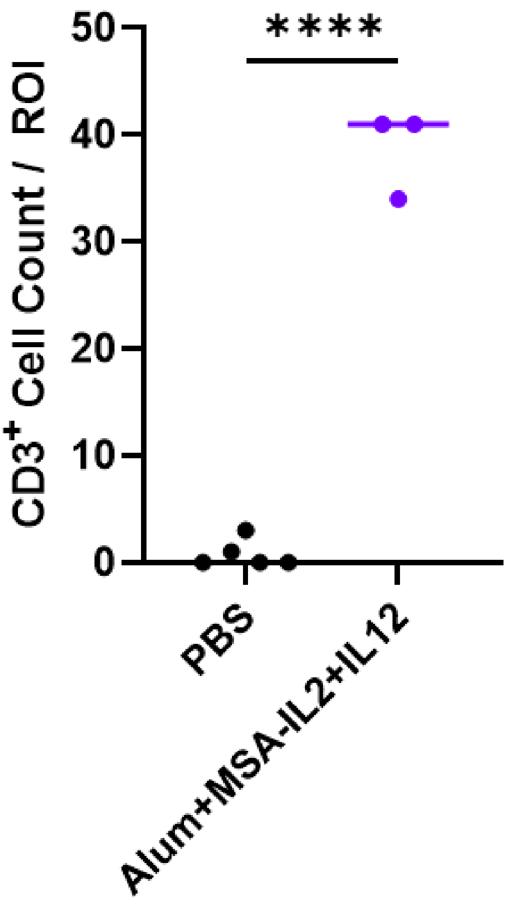


**Figure S3**: Quantification of CD3+ cells after administration of the therapeutic alum/MSA-IL2/IL12 depot. An unpaired t-test was used for statistical analysis. ****p<0.001.

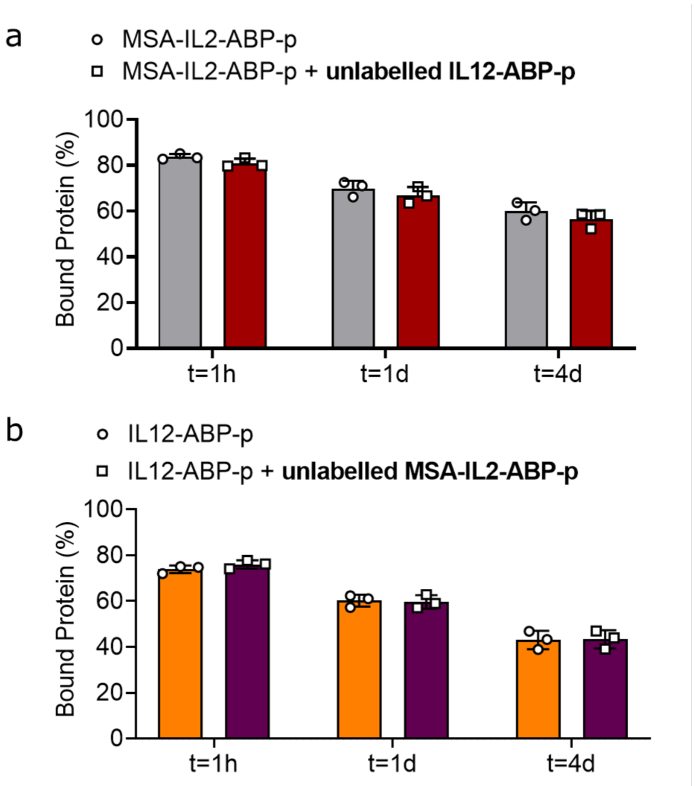


**Figure S4**: *In vitro* assay at various timepoints quantifying displacement and showing that binding of one protein does not displace the other on alum.

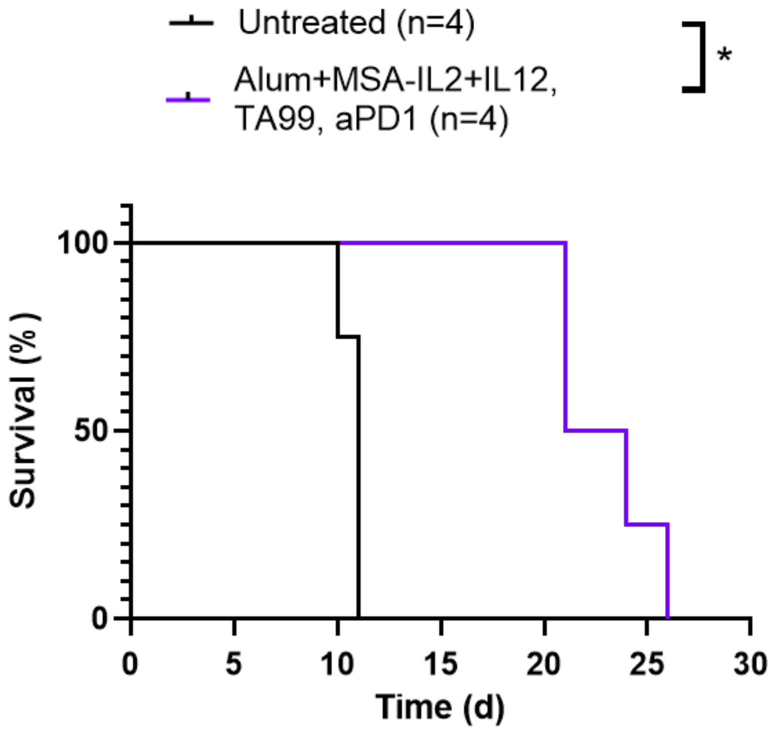


**Figure S5**: Survival curve of mice treated with alum/MSA-IL2/IL12 in the pre-motor cortex (M2). Statistical analysis was conducted with a Gehan-Breslow-Wilcoxon test. *p=0.0114.

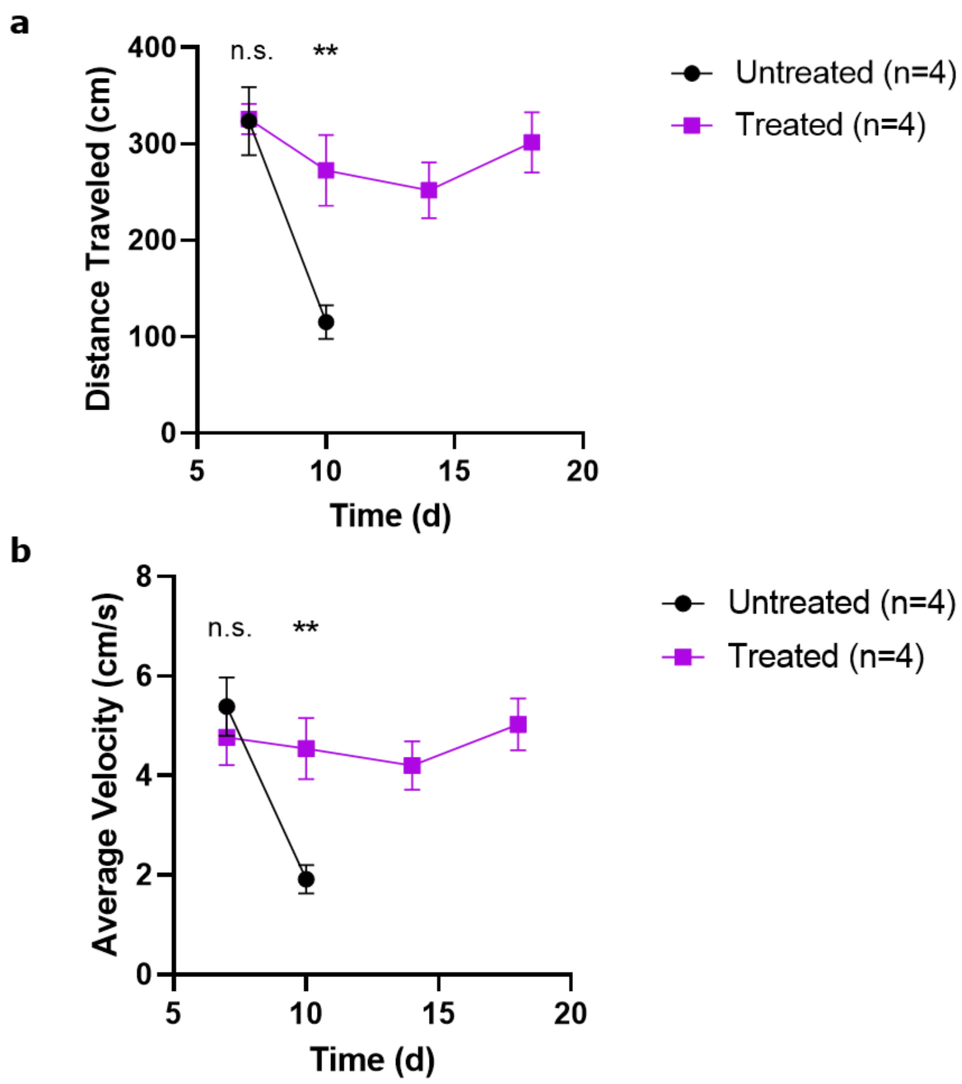


**Figure S6**: Open-field test on mice with B16F10 pre-motor cortex (M2) tumors. (a) Analysis of the average distance traveled during the open-field test. Statistical analysis was done on each day there were surviving mice in all groups with an unpaired t test. **p=0.0082. (b) Analysis of the average velocity during the open-field test. Statistical analysis was done on each day there were surviving mice in all groups with an unpaired t test. **p=0.0083.

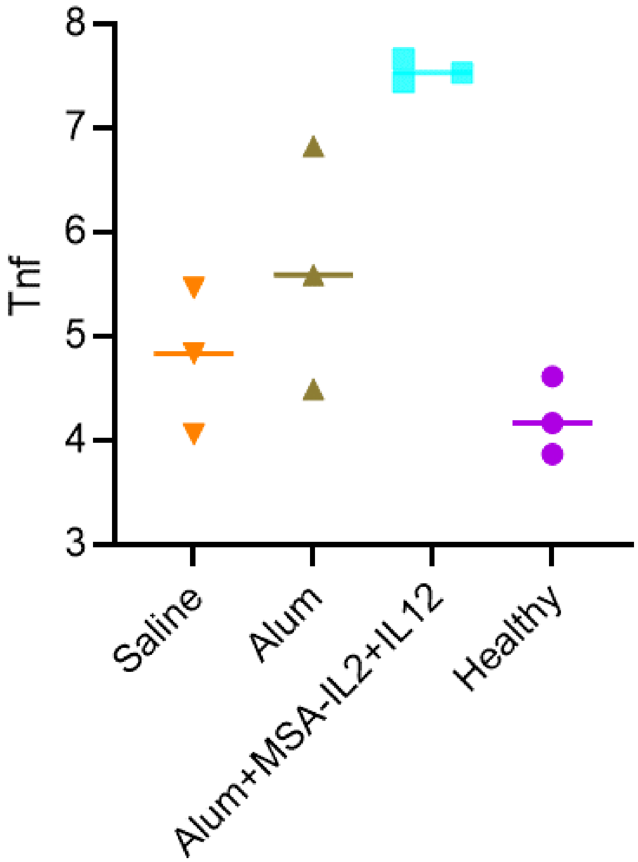


**Figure S7**: We performed RNA profiling using NanoString to obtain a more holistic immune profile of melanoma-bearing mice treated with saline, alum, or alum-tethered MSA-IL2 and IL12. Transcripts of *tnf* are plotted as $\log_2$ of normalized expression.

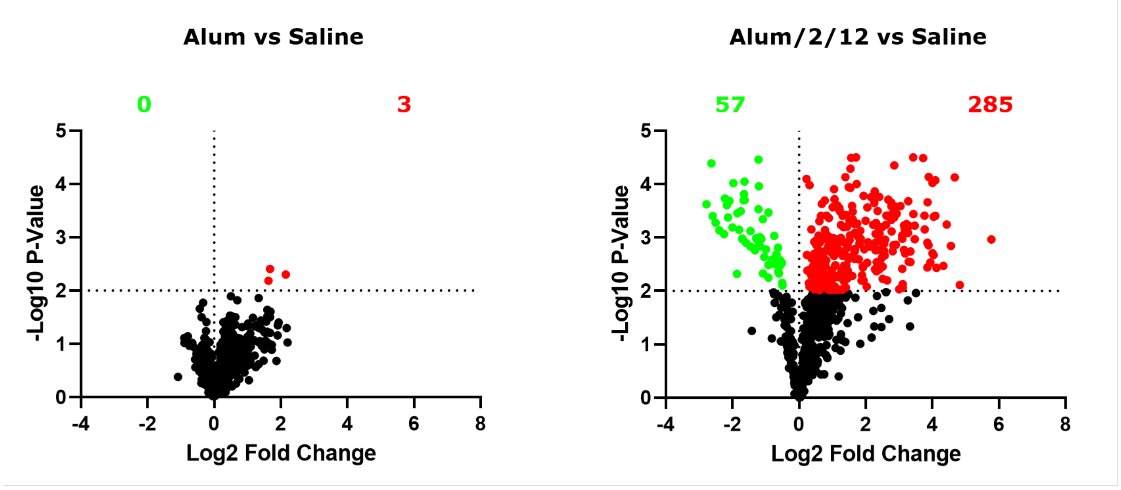


**Figure S8**: We performed RNA profiling using NanoString to obtain a more holistic immune profile of melanoma-bearing mice treated with saline vs. alum (left), or saline vs. alum with tethered MSA-IL2 and IL12 (right). Data shows 342 genes are differentially expressed (of 757 genes total) over control for alum/MSA-IL2/IL12, compared to only 3 genes for control for alum.

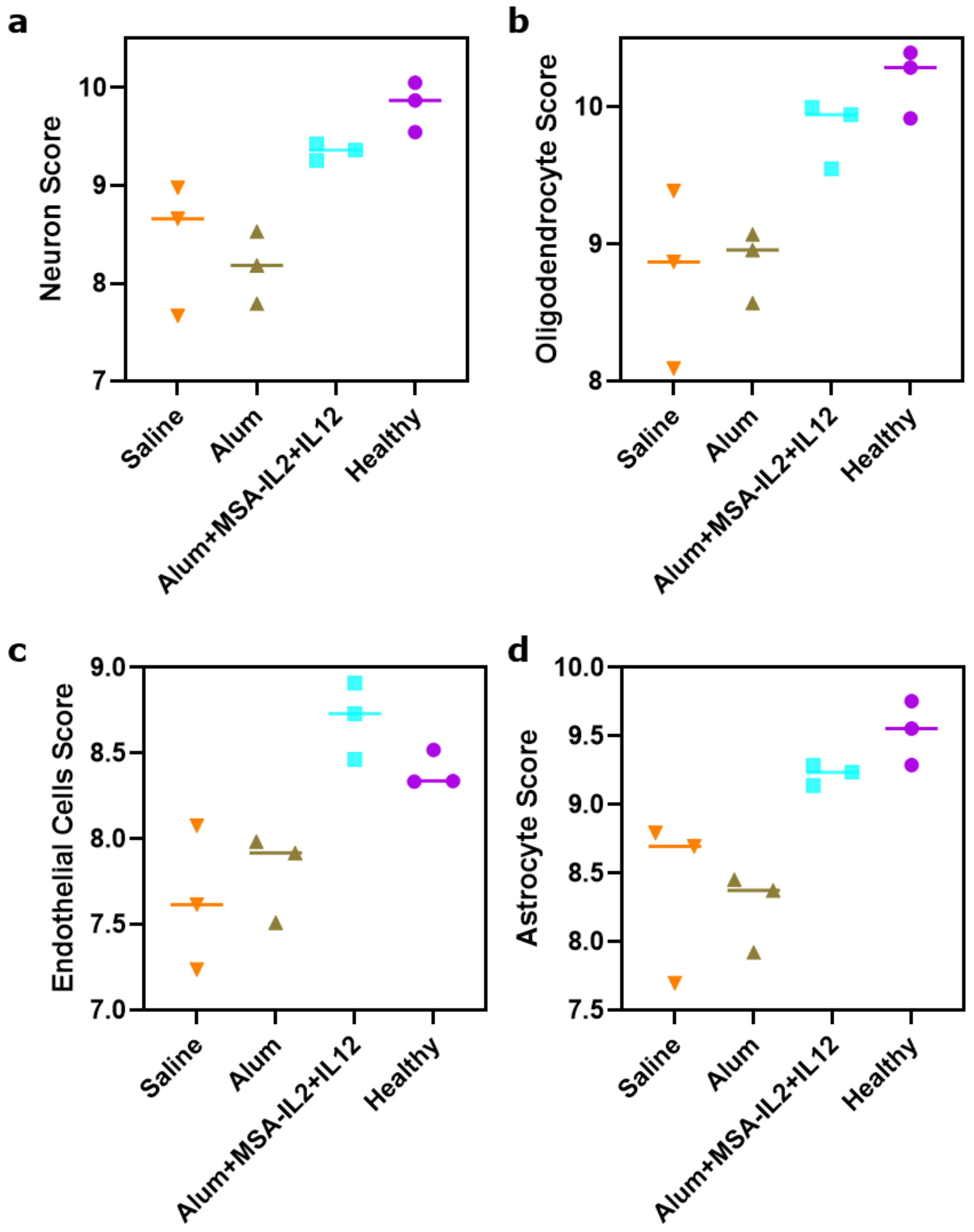


**Figure S9**: We performed RNA profiling using NanoString to obtain a more holistic neuronal profile of melanoma-bearing mice treated with saline, alum, or alum-tethered MSA-IL2 and IL12. Transcripts associated with (a) neurons, (b) oligodendrocytes, (c) endothelial cells, and (d) astrocytes are plotted. Data plotted is raw cell type abundance (NanoString) shown on a $\log_2$ scale, and is the average of the counts of characteristic genes associated with the cell type.

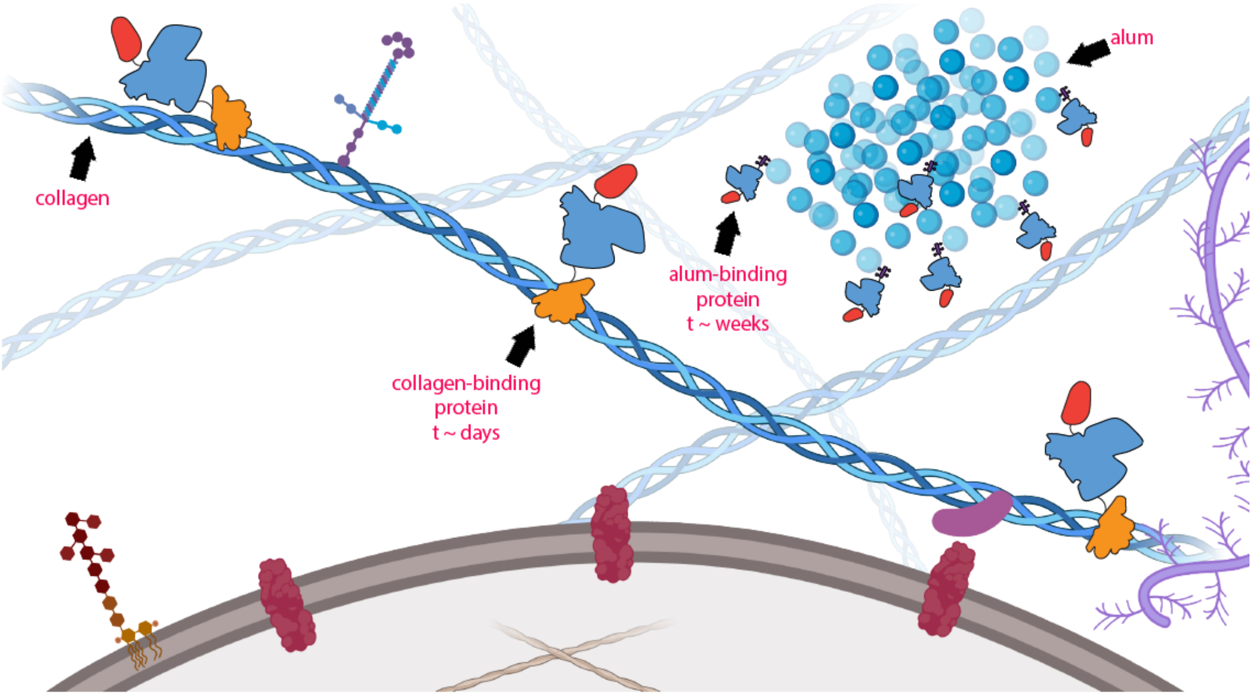


**Figure S10**: Graphical illustration demonstrating various engineered cytokine systems in the brain. Collagen-binding proteins persist on the timescale of days, whereas alum-tethered proteins persist on the timescale of weeks. Depending on the neuroimmune application, the duration and location of inflammation can be tuned. Figure made with Biorender.

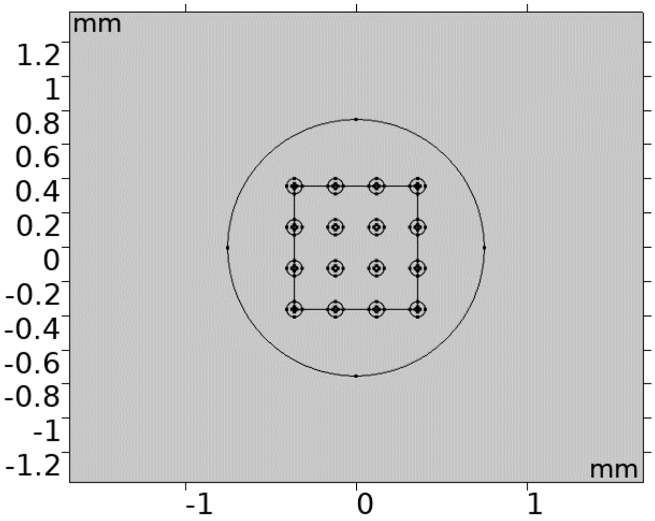


**Figure S11**: Model geometry used in Comsol to model relative protein retention. Model was first made in Solidworks 2021 and imported into Comsol.

### References


(1) Y. Agarwal, L. E. Milling, J. Y. H. Chang, L. Santollani, A. Sheen, E. A. Lutz, A. Tabet, J. Stinson, K. Ni, K. A. Rodrigues, T. J. Moyer, M. B. Melo, D. J. Irvine and K. D. Wittrup, *Nature Biomedical Engineering*, 2022, **6**, 129–143.

(2) N. Momin, J. R. Palmeri, E. A. Lutz, N. Jailkhani, H. Mak, A. Tabet, M. M. Chinn, B. H. Kang, V. Spanoudaki, R. O. Hynes and K. D. Wittrup, *Nature Communications*, 2022, **13**, 109.

(3) L. Ray, J. J. Iliff and J. J. Heys, *Fluids and Barriers of the CNS*, 2019, **16**, DOI: 10.1186/s12987-019-0126-9.

(4) M. M. Schmidt and K. D. Wittrup, *Molecular Cancer Therapeutics*, 2009, **8**, DOI: 10.1158/1535-7163.MCT-09-0195.

(5) G. M. Thurber and K. D. Wittrup, *Journal of Theoretical Biology*, 2012, **314**, DOI: 10.1016/j.jtbi.2012.08.034.

(6) K. D. Wittrup, G. M. Thurber, M. M. Schmidt and J. J. Rhoden, *Methods in Enzymology*, 2012, **503**, DOI: 10.1016/B978-0-12-396962-0.00010-0.

(7) Y. I. Wang, H. E. Abaci and M. L. Shuler, *Biotechnology and Bioengineering*, 2017, **114**, 184–194.

(8) N. Momin, N. K. Mehta, N. R. Bennett, L. Ma, J. R. Palmeri, M. M. Chinn, E. A. Lutz, B. Kang, D. J. Irvine, S. Spranger and K. D. Wittrup, *Science Translational Medicine*, 2019, **11**, DOI: 10.1126/scitranslmed.aaw2614.

(9) G. Paxinos and K. B. Franklin, *The Mouse Brain in Stereotaxic Coordinates, 2nd edition*, 2004.
